# Controlling Reciprocity in Binary and Weighted Networks: A Novel Density-Conserving Approach

**DOI:** 10.1101/2024.11.24.625064

**Authors:** Fatemeh Hadaeghi, Kayson Fakhar, Claus C. Hilgetag

## Abstract

We introduce efficient Network Reciprocity Control (NRC) algorithms for steering the degree of asymmetry and reciprocity in binary and weighted networks while preserving fundamental network properties. Our methods maintain edge density in binary networks and cumulative edge weight in weighted graphs. We test these algorithms on synthetic benchmark networks, including random, small-world, and modular structures, as well as brain connectivity maps (connectomes) from various species. We demonstrate how adjusting the asymmetry-reciprocity balance under edge density and total weight constraints influences key network features, including spectral properties, degree distributions, community structure, clustering, and path lengths. Additionally, we present a case study on the computational implications of graded reciprocity by solving a memory task within the reservoir computing framework. Furthermore, we establish the scalability of the NRC algorithms by applying them to networks of increasing size. These approaches enable systematic investigation of the relationship between directional asymmetry and network topology, with potential applications in computational and network sciences, social network analysis, and other fields studying complex network systems where the directionality of connections is essential.

## 1 Main

The intricate balance between reciprocity and asymmetry in complex networks is fundamental to the behavior of diverse systems, ranging from neural circuits to quantum many-body systems. Reciprocity, classically defined as the proportion of directed links with a reciprocal counterpart, is not merely a structural property, but a key factor influencing system-wide behaviors (Boguñá and Serrano, 2005; Garlaschelli and Loffredo, 2006). The balance of directed links profoundly impacts information flow, synchronization, and phase transitions across various network types (Pecora et al., 2014; Allard et al., 2024). Moreover, the degree of reciprocity or asymmetry in network connections plays a crucial role in determining the robustness and stability of real-world networks, governing critical phenomena observed in both natural and engineered systems (Rutishauser et al., 2015; Medeiros et al., 2021).

Despite its critical role, and for analytical convenience, the directed nature of connections is often simplified or overlooked in network models, neglecting the effects of varying levels of reciprocity and asymmetry.

However, directionality in real-world networks has a profound impact on their behavior. In directed networks, asymmetry introduces topological features such as trophic coherence (Johnson, 2020) and non-normality (Asllani et al., 2018), which significantly shape network dynamics. These structural properties manifest in the graphs’ eigenspectra, directly linking them to the stability and behavior of complex systems, including food webs, neuronal circuits, communication networks, and ecological systems (Zamora-López et al., 2008b; Johnson, 2020; Johnson et al., 2014; Ocklenburg and Mundorf, 2022). Moreover, structural asymmetry and network directionality have been identified as crucial factors in facilitating converse symmetry breaking or asymmetry-induced symmetry. These phenomena demonstrate how asymmetry, often viewed as a destabilizing feature, can paradoxically enable the stabilization of symmetric dynamical states, such as synchronization, in systems where fully symmetric coupling would otherwise fail (Nishikawa and Motter, 2016; Hart et al., 2019; Medeiros et al., 2021).

Reciprocal connections play an equally crucial role, significantly affecting dynamical processes, network growth, and the formation of higher-order structures such as motifs and communities (Boguñá and Serrano, 2005; Zlatic and Štefancic, 2011; Xu et al., 2022; Zamora-López et al., 2008b; Zlatić and Štefančić, 2009). In networks that transport information or materials, such as email systems, the World Wide Web, or social media platforms, bidirectional links are fundamental drivers of network dynamics, facilitating robust mutual exchange and enhancing the efficiency and reach of propagation processes (Zlatić and Štefančić, 2009; Zhu et al., 2014). This significance extends to other network types as well. In biological networks modeling dynamic processes, such as gene regulatory networks or neural circuits, reciprocal connections often represent feedback loops crucial for homeostasis and adaptive responses (Martinez et al., 2008; Ma et al., 2009). In social networks, bidirectional links can signify mutual trust or collaboration, influencing opinion formation and social cohesion (Zhu et al., 2014; Xu et al., 2022), while in economic networks, reciprocal trade relationships can impact market stability and economic resilience (Yao et al., 2021). Across these varied contexts, reciprocity enables immediate feedback, creating pathways for rapid signal propagation, resource sharing, and the emergence of complex behaviors, potentially leading to phenomena such as synchronization in coupled oscillators, pattern formation in reaction-diffusion systems, or cascading effects in financial networks.

Despite growing recognition of the importance of directionality and reciprocity in shaping network dynamics, studies have traditionally focused on extreme cases of complete symmetry or asymmetry, while the systematic exploration of intermediate levels has only recently gained attention. The current emphasis on extremes leaves a critical gap in understanding how the balance between reciprocity and asymmetry is optimized in real-world networks, as well as computational models such as artificial neural networks. While it is clear that the link balance significantly influences stability, functionality, and information flow, models capable of systematically manipulating reciprocity while preserving other key network properties remain limited; only a few notable examples have been proposed (Garlaschelli and Loffredo, 2006; Kasyanov et al., 2023; Allard et al., 2024). Here, we address this gap by introducing novel algorithms that precisely tune reciprocity in both binary and weighted networks, offering a new approach for directly exploring how reciprocity influences emergent behaviors in complex systems.

We demonstrate the broad applicability of our approach by analyzing how changes in reciprocity impact critical topological features of a network, including in- and out-degree distributions, spectral properties, modularity, and clustering coefficients. Additionally, we present a case study on the computational implications of graded reciprocity by solving a memory task (Damicelli et al., 2022) within the framework of reservoir computing (Jaeger and Haas, 2004). This application allows us to explore how varying levels of reciprocity affect a network’s ability to store and retrieve information, providing further insight into the broader functional consequences of reciprocity in both structural and dynamical contexts. Furthermore, we establish the scalability of our method across networks of varying sizes and initial reciprocity levels, ensuring its utility for real-world, large-scale network analyses.

## 2 Defining reciprocity in networks

In a directed binary network, represented by an adjacency matrix, *A*, where *A*_*ij*_ ∈ {0, 1}, let *L* denote the total number of links in the network, and 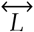 the number of reciprocal links, where a reciprocal link refers to a pair of directed edges (*i* → *j*) and (*i* → *j*). The link reciprocity is then quantified as the fraction of reciprocal links relative to the total number of directed links:

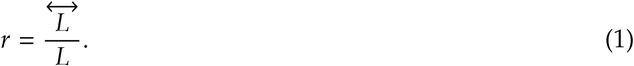

This definition of reciprocity, however, has been criticized for having only relative significance, as the measured value, *r*, must be compared to the expected reciprocity of a random graph with the same size and density. This comparison complicates the ability to rank different networks in terms of reciprocity, especially when comparing networks of varying density, because denser networks naturally exhibit more mutual connections by chance (Garlaschelli and Loffredo, 2004). We are aware of this criticism, but since our work aims to adjust the reciprocity of a given network while preserving its size and density (see 3 for details), our baseline remains the original network. In this context, it is acceptable for the adjusted reciprocity value to have a relative meaning, as it serves the purpose of measuring changes within a controlled framework.

Weighted networks, with the adjacency matrix *W* ∈ ℝ^*N*×*N*^, generalize unweighted (binary) networks by allowing links to have different real-valued weights, which can be positive, negative, or zero. The graph can be either undirected, where *W*_*ij*_ = *W*_*ji*_, or directed, where *W*_*ij*_ need not equal *W*_*ji*_.

While the reciprocity of binary networks has been studied extensively, the reciprocity of weighted networks has received much less attention. This fact is due to the more complicated phenomenology at the dyadic level in weighted networks. In a binary graph, it is straightforward to define link reciprocation - a link from node *i* to node *j* is reciprocated if the link from *j* to *i* also exists. However, in a weighted network, assessing whether a link of weight *W*_*ij*_ > 0 from node *i* to node *j* is reciprocated based on the weight *W*_*ji*_ of the mutual link is not trivial. Consequently, quantifying reciprocity in weighted networks poses additional challenges (Squartini et al., 2013).

Here, we designed our algorithms based on the formal definition of reciprocity in weighted networks proposed by Squartini et al. (Squartini et al., 2013). In this formalism, reciprocated weight between *i* and *j* (the symmetric component) is represented as

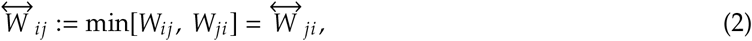

and the non-reciprocated weight from *i* to *j* (the asymmetric component) as

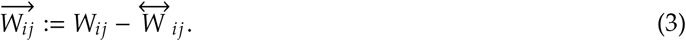

The strength reciprocity of a weighted network can, therefore, be computed as

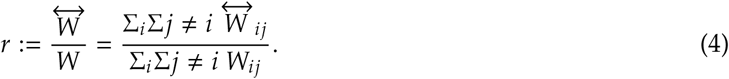

This quantity is, however, scale-invariant, such that if all the weights are multiplied by a scale factor, *W*_*ij*_ → *αW*_*ij*_, *r* does not change, as shown in (Squartini et al., 2013). This aspect makes the task of adjusting strength reciprocity challenging, particularly under the constraint of preserving cumulative edge weights, *E* = Σ*W*_*ij*_.

## 3 Novel algorithms for modulating network reciprocity

The proposed Network Reciprocity Control (NRC) algorithms operate under several key assumptions. First, they assume that unidirectional links have the highest potential to become reciprocal when adjusting the network towards the desired reciprocity, reflecting the idea that existing connections may be easily converted into bidirectional ones. Additionally, the algorithms presume that converting reciprocal links into unidirectional links does not significantly disrupt the overall network structure, allowing for flexible reciprocity adjustments. Throughout the process, the network density in binary graphs and the total strength in weighted networks are maintained. In the binary case, the network is treated as maximally random while adhering to its specified link density and desired reciprocity, thereby minimizing systematic biases during the adjustment process. Unlike methods that fix both in- and out-degree sequences (Garlaschelli and Loffredo, 2006; Kasyanov et al., 2023; Allard et al., 2024; Zamora-López et al., 2008a), our approach does not constrain degree correlations. This design choice enables a broader exploration of how reciprocity influences network structure and function, although it can also permit changes in topological features such as degree correlations or clustering. However, our analyses, presented in section 5, show that the algorithm preserves the average in-degree and out-degree, and that changes in clustering coefficient are generally small across the tested networks. This balance provides a flexible framework for investigating reciprocity, complementary to more constrained models that strictly control multiple structural parameters.

The algorithm designed to adjust reciprocity in the weighted network uses a deterministic strategy. Both algorithms assume symmetric treatment of link or weight modifications, meaning that changes are balanced to preserve the network’s topology. Local adjustments in link or weight configurations are considered to have global effects on the network’s reciprocity, and the algorithms are designed to approximate the target reciprocity within a small tolerance interval, acknowledging that perfect alignment with the desired value may not always be feasible.

### 3.1 Binary networks

#### 3.1.1 Algorithm design and implementation

Given a directed binary network represented by an adjacency matrix, *A*, where *A*_*ij*_ ∈ { 0, 1 }, the algorithm, detailed in **Algorithm 1 (binary)**, aims to adjust the network’s reciprocity to a specified target value, *r*_*d*_ while maintaining the overall structure and the number of links, i.e, *L* = Σ_*i*≠*j*_ *A*_*ij*_.

##### Step 1: Initial setup

The current reciprocity, *r*_*c*_, is calculated as the ratio of reciprocal links to the total number of links. If *r*_*c*_ is already within a small tolerance interval of the desired reciprocity, *r*_*d*_, the algorithm terminates without making changes.

##### Step 2: Identification of symmetric and asymmetric links

Symmetric (reciprocal) links, denoted by *S*, are identified where *A*_*ij*_ = 1 and *A*_*ji*_ = 1, with *i* < *j* to prevent double counting. Asymmetric 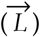 links are identified where *A*_*ij*_ = 1 and *A*_*ji*_ = 0.

##### Step 3: Determine target number of reciprocal links

The target number of reciprocal links, 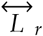, is estimated using

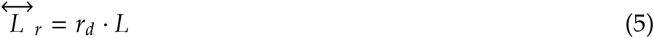

where *L* represents the total number of links.

##### Step 4: Reciprocity adjustment

- Case 1: Reducing reciprocity (*r*_*d*_ < *r*_*c*_) If the number of reciprocal links, | *S*|, is found to exceed the target value 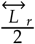, the difference is computed as

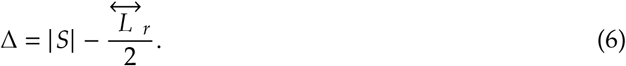 To reduce reciprocity, Δ reciprocal links are removed from *S* by converting them randomly into unidirectional links. To avoid introducing directional bias when reducing reciprocity, pruned links are selected with equal probability from the upper and lower triangles of the adjacency matrix. This ensures that no systematic preference is given to one direction over the other (i.e., *i* → *j* or *j* → *i*), maintaining the overall balance of directionality in the network structure. To maintain network density, Δ unidirectional links are distributed to Δ non-existing connections, ensuring that they are not already reciprocal.
- Case 2: Increasing reciprocity (*r*_*d*_ > *r*_*c*_) When increasing reciprocity, the difference Δ, as computed in Eq. 6, indicates the number of ≥ unidirectional links that should be converted to reciprocal links. If sufficient unidirectional links are available 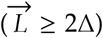, Δ asymmetric links are converted to reciprocal links to increase reciprocity, and Δ are removed from other unidirectional links to preserve density. If unidirectional links are insufficient 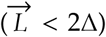, all available unidirectional links are converted to reciprocal links. Excess reciprocal links are then removed, and asymmetric links are added to randomly chosen non-existing connections to achieve the desired reciprocity while maintaining network density.

#### 3.1.2 Bounds of reciprocity: Theoretical limits and practical implications

For a directed binary network characterized by an adjacency matrix *A*, with the total number of links denoted as 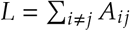 our algorithm does not impose constraints on the correlation between in-degrees and out-degrees. This flexibility allows the network to attain full reciprocity, where the reciprocity index *r* = 1.0. The lower bound of reciprocity, however, is determined by the size of the network and its link density. This section derives the mathematical lower bound of reciprocity for a network with *N* nodes. The total number of links, *L*, can be decomposed into two categories: reciprocal links 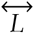, and asymmetric links, 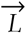, such that:

#### Algorithm 1

Adjust Reciprocity of Binary Network

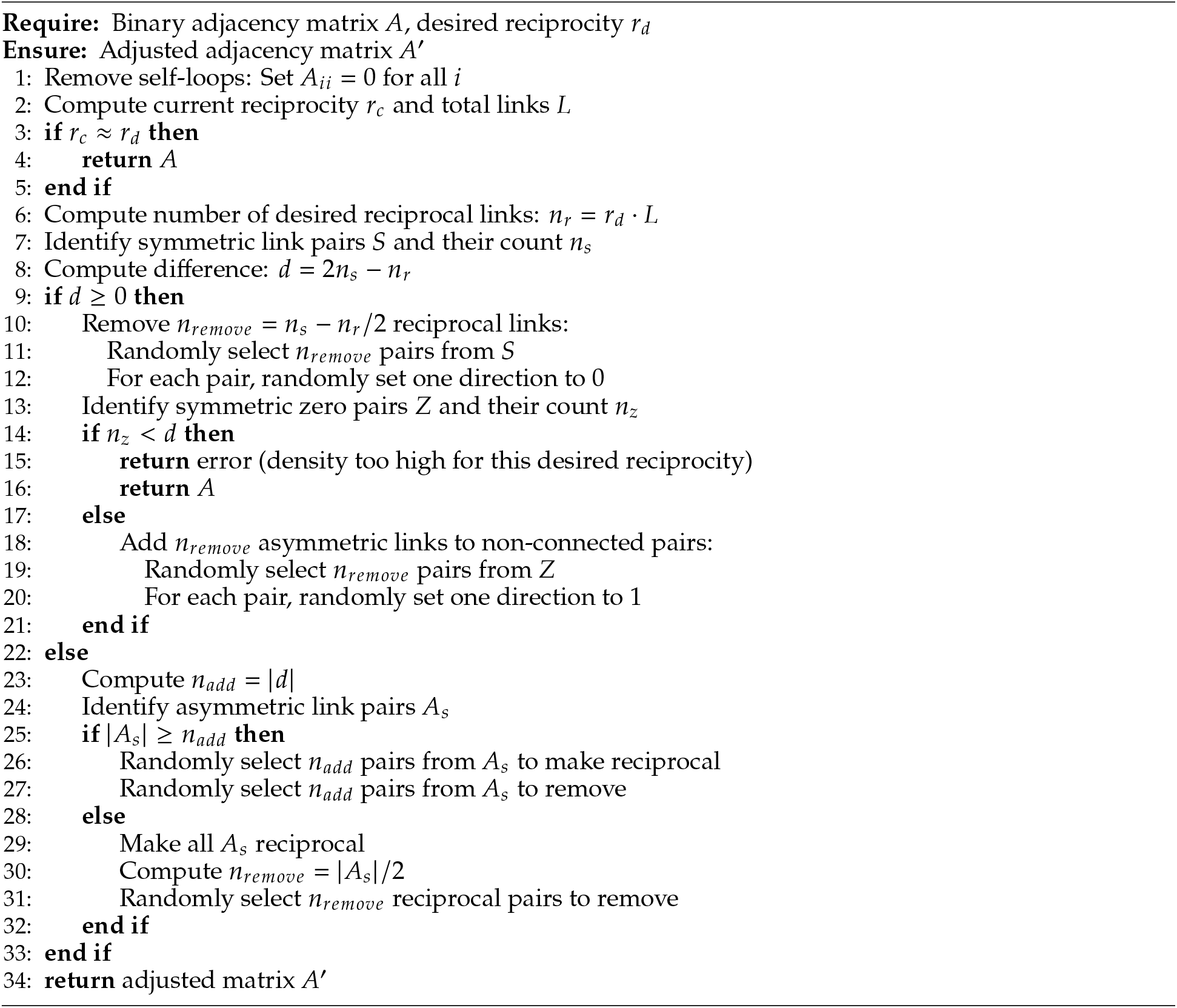

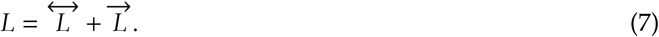

For a sparse network with 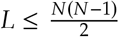, it is possible to generate a maximally random configuration of the adjacency matrix *A* where all links are asymmetric 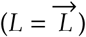 and no reciprocal pairs are present 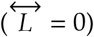. In this scenario, the lower bound of reciprocity is 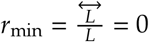.

However, for networks where 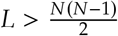, such that 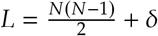 with *δ* > 0, at least 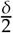 reciprocal link pairs must exist to maintain the link density. In this case, the lower bound of reciprocity is g iven by:

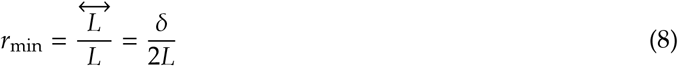

This formalism defines the bounds for asymmetry and shows how network size and density set a lower limit on the reciprocity achievable by our algorithm. Note that in the thermodynamic limit (*N* → ∞), a fixed additive surplus *δ* becomes negligible relative to L, and the minimum reciprocity approaches 0. Our formulation applies to finite, sparse networks, where 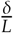 remains positive.

#### 3.1.3 Global connectivity during reciprocity adjustment

An important consideration when modifying reciprocity is the potential impact on global connectivity. Specifically, one might ask whether converting unidirectional links to reciprocal pairs, or pruning reciprocal links to create asymmetric connections, could fragment the network or disrupt the giant connected component. Although Algorithm 1 (binary) does not explicitly enforce preservation of a giant component during reciprocity adjustment, we consistently observed that the giant connected component remained intact across all tested networks, including random, small-world, and modular structures. In modular networks with very low inter-module link density, there is, in principle, a greater risk that pruning asymmetric inter-module links could reduce connectivity between modules. However, such fragmentation did not occur in our experiments. Future work could incorporate explicit constraints or post-adjustment validation steps to ensure the preservation of global connectivity where this is critical for specific applications.

### 3.2 Weighted networks

#### 3.2.1 Algorithm design and implementation

Given a weighted adjacency matrix *W* ∈ ℝ^*N*×*N*^, the following algorithm, detailed in **Algorithm 2 (we**I;**ighted)**, is proposed to adjust strength reciprocity while preserving the sum of connection strengths, *E* = should be noted that weights are restricted to non-negative values, i.e., *W*_*ij*_ ≥ 0 for *i* = 1, 2, …, *N*.

##### Step 1: Initial setup

The current reciprocity, *r*_*c*_, is computed using Eq. 4. If *r*_*c*_ is found to be close to the desired strength reciprocity, *r*_*d*_ (within a small tolerance), the algorithm is terminated without making any adjustments.

##### Step 2: Identification of symmetric and asymmetric network components

The symmetric component of the adjacency matrix, denoted by 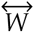, is identified where *W*_*ij*_ = *W*_*ji*_ = min[*W*_*ij*_, *W*_*ji*_] (see Eq. 2). The asymmetric component, 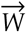, is then identified as the difference 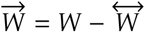.

##### Step 3: Determination of reciprocity scaling factor

The reciprocity scaling factor, *β*, is computed based on the desired strength reciprocity, *r*_*d*_, as:

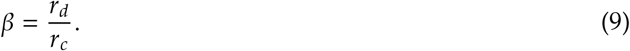

##### Step 4: Reciprocity adjustment

- Case 1: Reducing reciprocity (*β* ≤ 1)

If a reduction in strength reciprocity is required, the symmetric component of the adjacency matrix is scaled down by a factor of *β*, such that 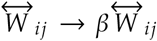. To preserve the total strength, *E*, the asymmetric component, 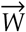, is scaled up accordingly. The decrease in the symmetric component reduces the total sum by:

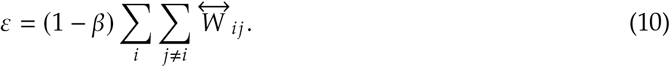

To compensate for this reduction, the positive elements of the asymmetric component are adjusted. First, the indices of positive elements in 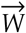 are identified as:

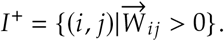

The number of such elements is counted as *n*^+^ = | *I*^+^ |. If *n*^+^ > 0, the increment to be added to each positive element is computed as:

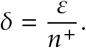

###### Algorithm 2

Adjust Strength Reciprocity While Preserving Total Connection Strengths

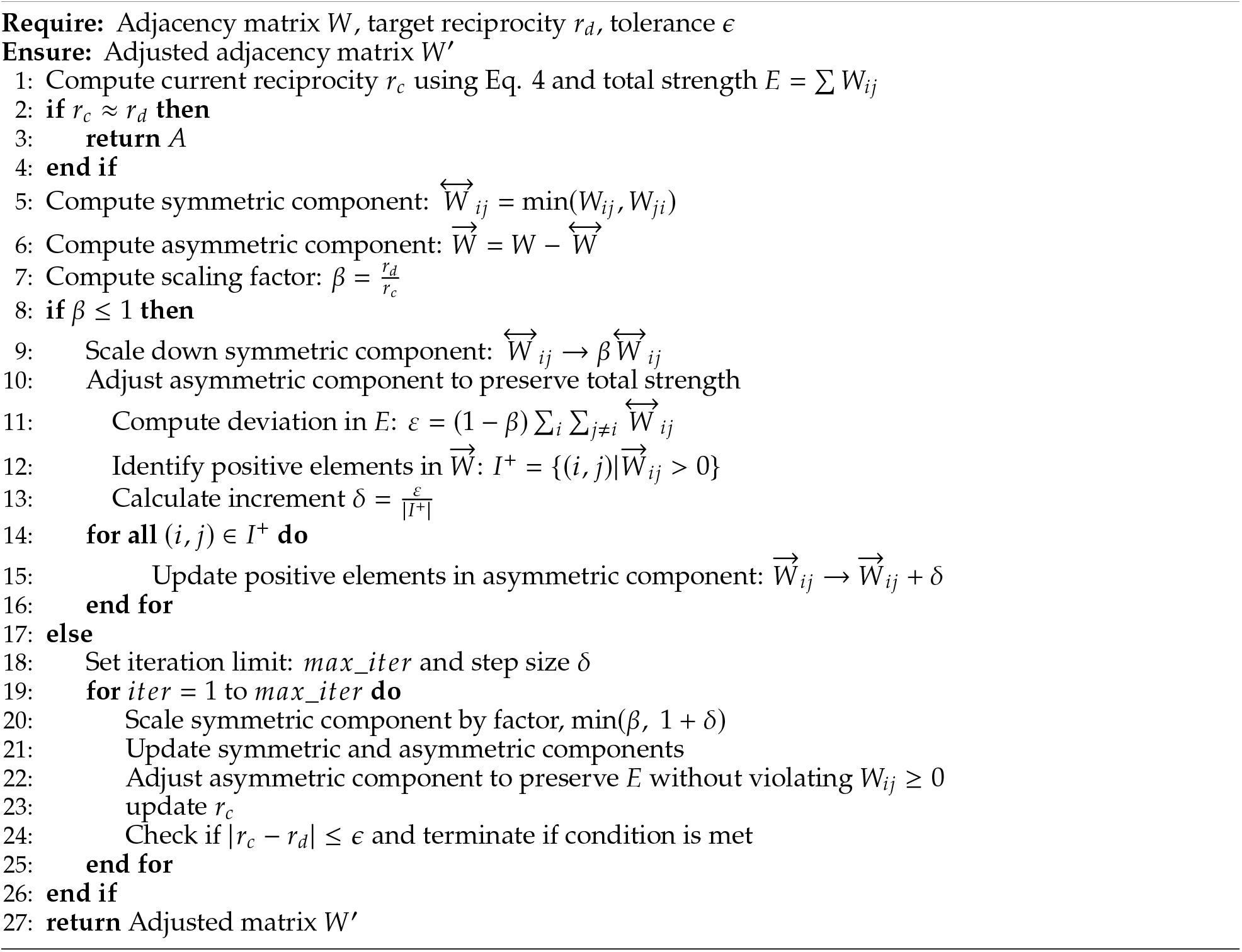

Finally, each positive element in 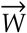 is updated as:

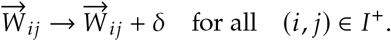

This procedure ensures that the total deviation, ε, is evenly distributed while maintaining the total weight sum and preserving the sign of each matrix element.

- Case 2: Increasing reciprocity (*β* > 1)

If an increase in strength reciprocity is needed, a direct scaling of the symmetric component is not feasible, as it may affect the values of min *W*_*ij*_, *W*_*ji*_ and alter the symmetric part of the matrix. Therefore, an iterative procedure is followed. At each iteration, the scaling factor is recalculated as the minimum of the ratio of the target to the current reciprocity, *β*, and *r*_*c*_ + *δ*, where *δ* is a small step size. This factor controls the gradual scaling of elements in 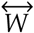. As the symmetric component is adjusted, the asymmetric component is reduced to maintain the total sum of strengths, *E*, ensuring that no values fall below a small, positive non-zero threshold to satisfy *W*_*ij*_ ≥ 0.

#### 3.2.2 Handling edge cases and exceptional scenarios

In this section, we elaborate on two extreme cases of network reciprocity: complete symmetry (*r* = 1.0) and complete asymmetry (*r* = 0), as handled by our algorithm. These cases allow us to examine how the network evolves under maximum structural constraints while preserving key properties such as total connection strength.

##### a. Symmetry Breaking in Initially Symmetric Networks

For weighted graphs *W* ∈ ℝ^*N*×*N*^ with initial reciprocity *r*_*c*_ = 1.0, we introduce a small perturbation to break structural symmetry. This is achieved by applying a random mask *P* ~ 𝒩 (0, *δ*), where *δ* is chosen to be smaller than the minimum non-zero element in *W*. This approach ensures that the perturbation is subtle enough to maintain the overall network structure while introducing the desired asymmetry.

##### b. Achieving Full Symmetry

While a fully symmetric network can be trivially obtained as 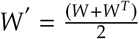, this method may introduce spurious connections, altering the network’s fundamental structure.^2^To maintain consistency in our analysis of network features (detailed in Section 5), we instead employ our iterative algorithm to adjust the strength reciprocity to *r*_*d*_ = 1.0. This approach preserves the original connection patterns while changing the strengths to achieve the desired symmetry.

##### c. Maximum Asymmetry

To explore the opposite extreme, we develop a method to create fully asymmetric networks. Given a weighted network *W* ℝ ∈ ^*N*×*N*^, we set all reciprocal weights to zero 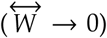 and up-scale the remaining non-zero elements to preserve the total edge weight *E*, while ensuring *W*_*ij*_ ≥ 0. It is important to note that this is the only scenario in which our algorithm removes existing links; in all other cases, unlike its binary counterpart, our method preserves the original network connectivity.

## 4 Experimental setup and data

The algorithms were applied to a range of synthetic networks, including random, small-world, and modular topologies, as well as empirical connectomes from multiple species. The analysis reveals how modifying the asymmetry-reciprocity balance impacts key network characteristics, including spectral properties, degree distributions, community structures, clustering coefficients, and shortest path lengths.

Each network type was generated over 50 independent trials using different random seeds to ensure distinct network realizations. Random networks were constructed as binary directed Erdős–Rényi graphs (Batagelj and Brandes, 2005). Small-world networks were modeled as connected Watts–Strogatz graphs (Watts and Strogatz, 1998), originally undirected, with a rewiring probability of 0.5 and neighborhood sizes *k* ∈ { 25, 26, 27, 28, 29 }, chosen to maintain an average density of approximately 0.1 while avoiding identical network samples. Modular networks were generated using a stochastic block model (Holland et al., 1983) with four equally sized blocks (64 nodes each) and symmetric connection probabilities defined by the matrix

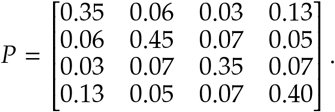

For weighted networks, random non-negative weights (*W*_*ij*_ > 0) sampled from a uniform distribution over [0, 1) were assigned to the existing links in these previously generated binary networks.

The algorithms were further tested on empirical connectomes from various species, which exhibit diverse sizes, densities, and initial binary or strength reciprocity values (Varshney et al., 2011; Chiang et al., 2011; Griffa et al., 2019; Modha and Singh, 2010; Rubinov et al., 2015; Bota et al., 2015). These key properties across datasets are summarized in Table 1. Representative synthetic and empirical connectomes are illustrated in Figure 1.

**Table 1.** Summary of synthetic and empirical networks analyzed in this study. The networks vary in size, density, and original reciprocity. For synthetic networks, different structures include random, small-world, and modular types. For a network with *N* nodes and *L* links, the density, *d*, was computed for directed networks as *L* /*N* (*N* ~1). For empirical networks, both weighted and binary representations are considered. The notation *r*_*w*_ denotes strength reciprocity, while *r*_*b*_ refers to binary reciprocity. For empirical networks originally weighted, a binary representation is generated by establishing a link where *W*_*ij*_ > 0.

| network | $N$ | $d$ | $r_w$ | $r_b$ |
| --- | --- | --- | --- | --- |
| **Synthetic networks** |  |  |  |  |
| Random | 256 | 0.05 | 0.032 | 0.07 |
| Small-world | 256 | 0.1 | 0.67 | 1.0 |
| Modular | 256 | 0.1 | 0.67 | 1.0 |
| **Connectomes** |  |  |  |  |
| C. elegans | 279 | 0.03 | 0.12 | 0.21 |
| Drosophila | 49 | 0.83 | 0.67 | 0.92 |
| Human-struct-scale033 | 83 | 0.31 | 1.0 | 1.0 |
| Human-struct-scale060 | 129 | 0.23 | 1.0 | 1.0 |
| Human-struct-scale125 | 234 | 0.13 | 1.0 | 1.0 |
| Human-struct-scale250 | 463 | 0.06 | 1.0 | 1.0 |
| Human-struct-scale500 | 1015 | 0.02 | 1.0 | 1.0 |
| Macaque* | 242 | 0.07 | – | 0.51 |
| Mouse | 112 | 0.53 | 0.52 | 0.67 |
| Rat | 73 | 0.37 | 0.45 | 0.66 |

\* Macaque connectome is only available as a binary network.

**Figure 1.**
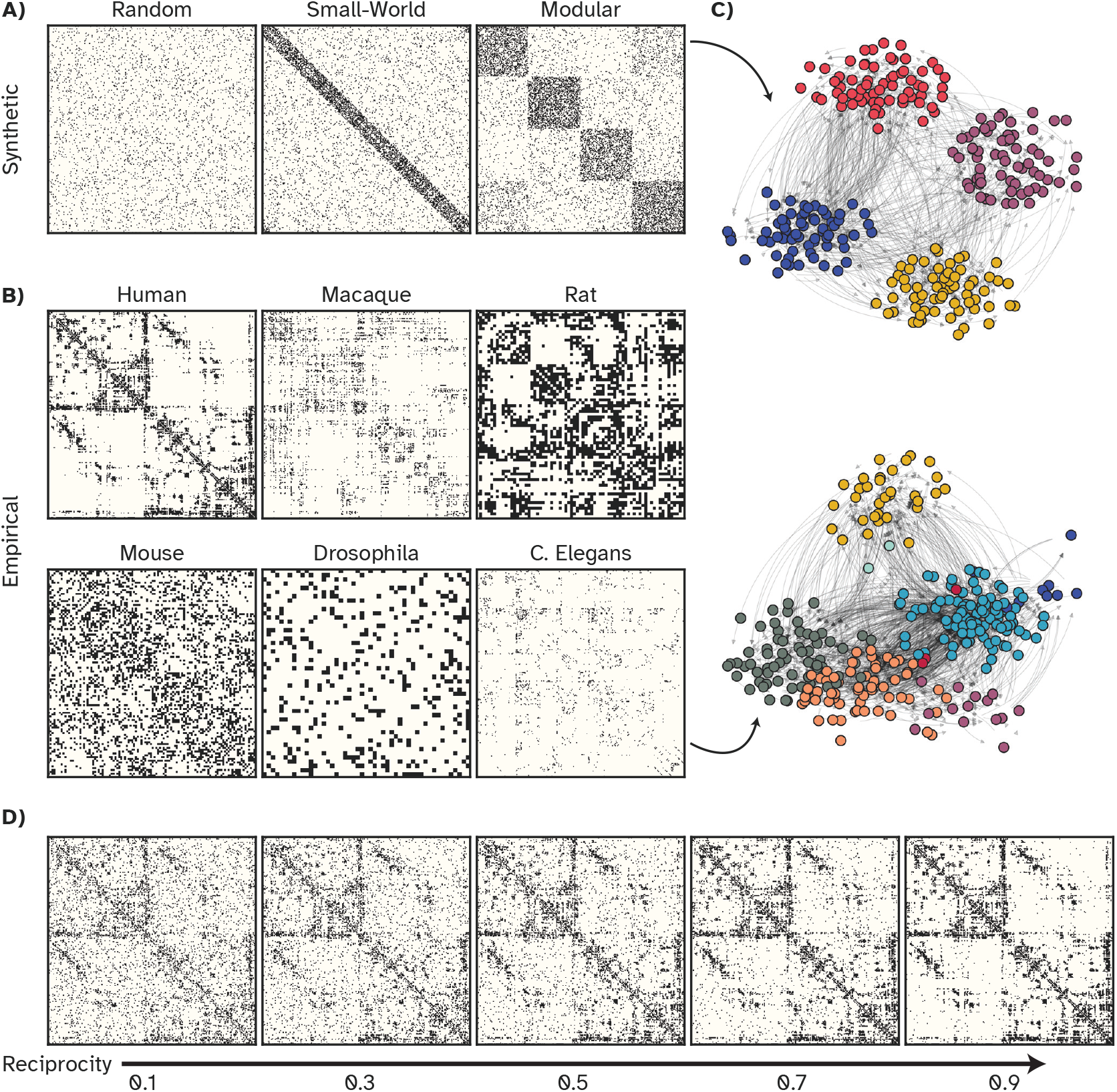
Representative synthetic (A) and empirical (B) connectomes used for algorithm testing. (A) The random network, initially with reciprocity *r*_*b*_ = 0.07, is shown after adjusting its reciprocity to *r* = 1.0. Small-world and modular networks are originally symmetric. Illustrated connectomes display the adjacency matrices after adjusting their reciprocity to *r*_*b*_ = 1.0 using Algorithm 1. (C) Community structures identified in two networks with clear modular organization. (D) Human connectome (Human-struct-scale125) with initial reciprocity, *r*_*b*_ = 1.0 (fully symmetric), shown alongside representations with adjusted reciprocity values, *r*_*b*_ = [0.1, 0.3, 0.5, 0.7, 0.9], from left to right.

## 5 Impact on network properties

### 5.1 Spectral analysis

This investigation reveals a clear connection between network reciprocity and the eigenvalue spectrum of the adjacency matrix, a critical measure of global network properties. Through systematic adjustment of reciprocity using the density-preserving algorithm, distinct modifications in the spectral characteristics of both random and structured networks are observed.

As illustrated in Figure 2, an increase in reciprocity is associated with a broadening of the distribution of eigenvalue magnitudes across 50 experimental trials. The symmetry of the real components of the eigenvalues is universally preserved. However, in structured networks, a subtle translation of the symmetry axis is detected when applying the binary algorithm. This phenomenon is absent in the weighted algorithm implementation shown in Figure B1.

**Figure 2.**
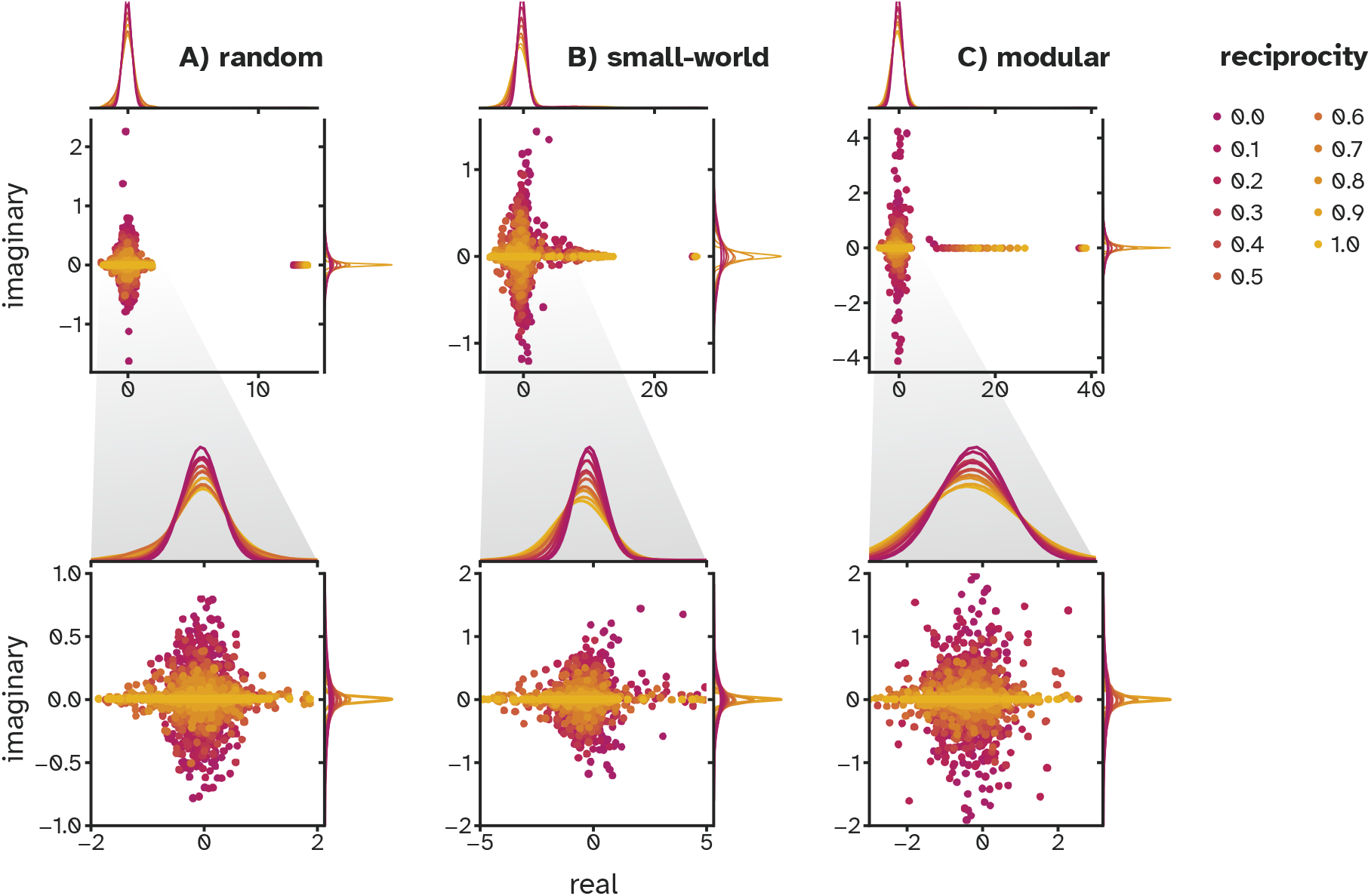
Functional implications of controlling reciprocity. Change in eigenspectrum of adjacency matrices as their reciprocity is adjusted from *r*_*b*_ = 0 to *r*_*b*_ = 1.0 using Algorithm 1 (binary). The top panel shows how eigenvalues are arranged in the complex plane, while the bottom panel provides a zoomed-in view. Distributions of real and imaginary components at various degrees of reciprocity were calculated from data collected in 50 trials for each setting.

To quantify the impact of reciprocity on matrix normality, Henrici’s index of departure from normality was employed (Asllani et al., 2018). This index, detailed in Appendix Non-normality, reaches its minimum for normal matrices and increases as matrices deviate from normality. The findings demonstrate that elevated reciprocity consistently enhances normality. This observation holds for both binary and weighted algorithmic approaches, as shown in Figure A1, and aligns with the results reported previously (Asllani et al., 2018).

These results provide crucial insights into the interplay between local dyadic relationships and global network topology, unencumbered by confounding density effects. The ability to fine-tune reciprocity while maintaining network density opens new avenues for investigating dynamical processes on directed networks.

### 5.2 Degree distribution

The algorithm for adjusting network reciprocity presents a novel approach compared to previous methods that operate under prescribed degree correlation constraints (Zamora-López et al., 2008a; Allard et al., 2024). By relaxing these constraints, a more comprehensive exploration of how correlation patterns evolve with changing reciprocity values is enabled.

As expected, fully symmetric networks (*r* = 1.0) exhibit perfect in-/out-degree correlation due to structural constraints. More interestingly, we observe that as reciprocity decreases, the degree correlation weakens, though the extent of the decline varies across network topologies. As shown in Figure 3, the reduction is more pronounced in random graphs and less evident in structured networks, underscoring the moderating effect of topology on reciprocity-driven changes in degree correlations.

**Figure 3.**
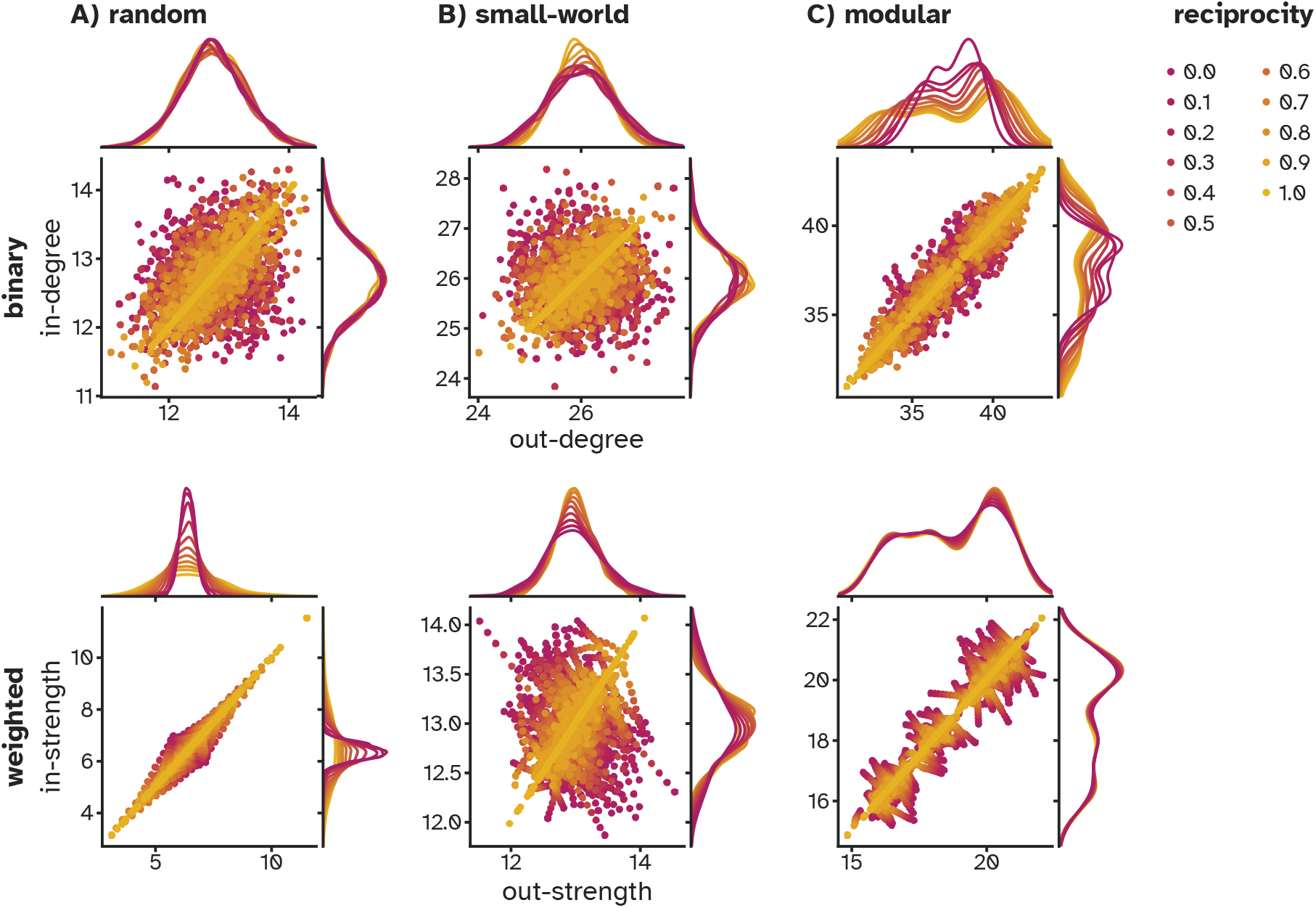
Structural implications of controlling reciprocity. Evolution of degree distributions and correlation in binary (top) and weighted (bottom) adjacency matrices as their reciprocity is adjusted from *r* = 0 to *r* = 1.0 using Algorithm 1 (binary) and Algorithm 2 (weighted).

Interestingly, while reciprocity variations in random networks do not alter the distributions of in- and out-degrees, structured networks exhibit subtle but important changes. In small-world networks, starting from initial reciprocity of 1.0, we observe a slight broadening of both in- and out-degree distributions as unidirectional links increase. Conversely, networks with modular structures display an opposite trend: the introduction of asymmetric links leads to a sharpening of both distributions, despite preserving their overall shape.

In weighted networks, the sensitivity of in- and out-strength distributions to reciprocity adjustments appears to be more topology-dependent. Networks with modular structures demonstrate more robust distributions, while random networks exhibit greater susceptibility to changes. Specifically, in random weighted networks, adjusting reciprocity to higher values (i.e., *r*_*d*_ > *r*_*c*_ = 0.07) leads to a noticeable broadening of both in- and out-strength distributions, reflecting the lack of inherent structural constraints that might otherwise stabilize these distributions against perturbations.

These findings illuminate the intricate interplay between reciprocity, network structure, and degree correlations. The development of an algorithm that enables systematic manipulation of reciprocity provides a powerful tool for comprehensive numerical exploration of these relationships.

### 5.3 Alterations in path lengths and network efficiency

The impact of reciprocity variation on inter-nodal distances within the network models was investigated. This analysis involved the transformation of weight matrices into length matrices through normalization and inversion of non-zero weights, followed by the application of the Floyd-Warshall algorithm to compute shortest paths. From these computations, global efficiency metrics (Seguin et al., 2018; Latora and Marchiori, 2001) were derived by averaging inverse shortest paths, and changes in network diameters were examined.

In binary networks, the results, depicted in Figure 4, reveal a striking resilience of these metrics to reciprocity alterations, with one notable exception: the *C. elegan*s connectome. Due to its extreme sparsity and initial reciprocity of 0.21, both increases and decreases in reciprocity led to significant enhancements in global efficiency.

**Figure 4.**
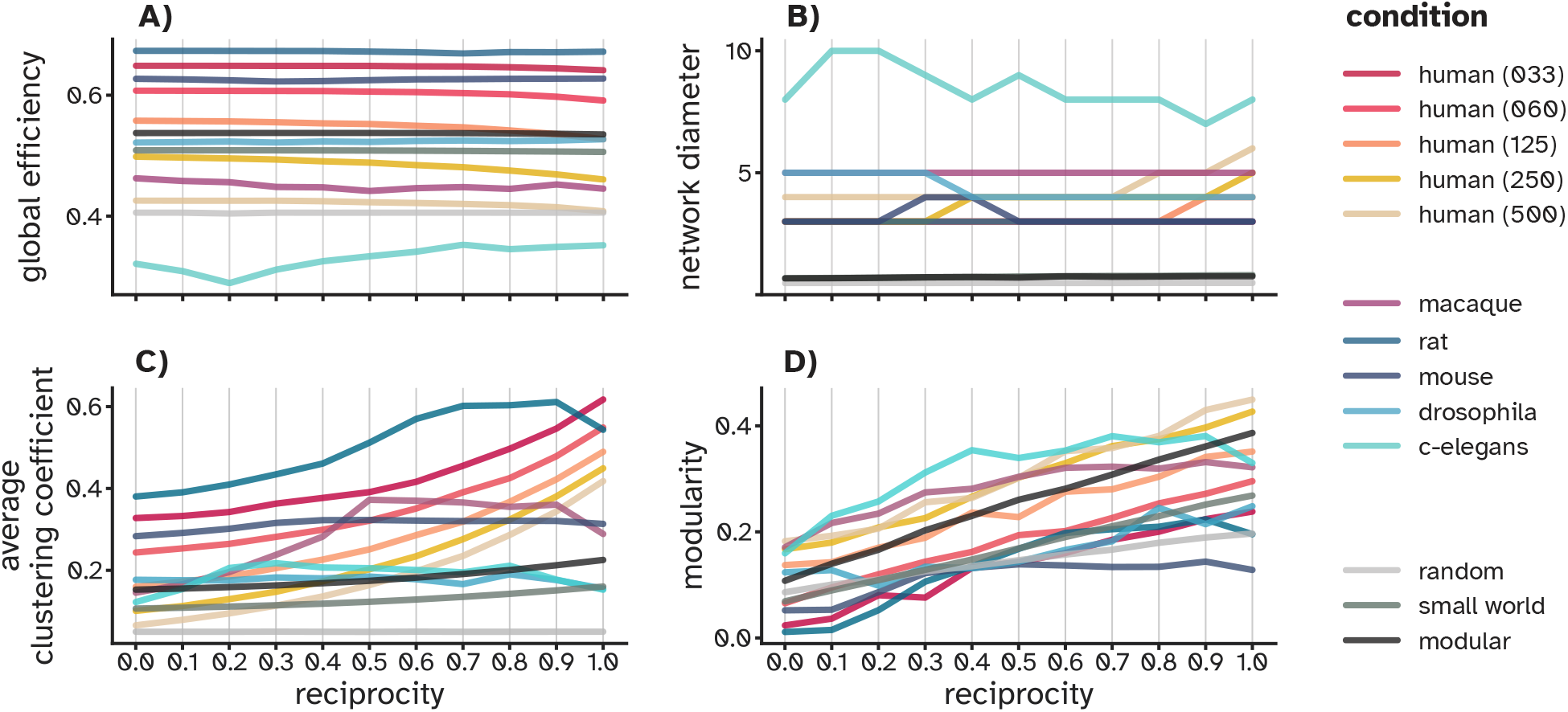
Structural implications of controlling reciprocity. Trends in key network metrics—global efficiency, network diameter, modularity index, and average clustering coefficient—in binary adjacency matrices as their reciprocity is adjusted from *r* = 0 to *r* = 1.0 using Algorithm 1 (binary).

Upon application of the weighted variant of the algorithm, an increase in global efficiency was observed as the asymmetric components of human connectomes were enhanced, as shown in Figure B2. This effect was not limited to biological networks, but was also observed in randomly generated network structures.

### 5.4 Evolution of community structure

In this analysis, the influence of reciprocity on network community structure was investigated using the modularity index (Clauset et al., 2004), detailed in Modularity index in the Appendix, as the primary metric. As shown in Figure 4, the modularity index increases consistently with rising network reciprocity, particularly in networks with an initially modular structure. The modular networks were initially constructed with complete symmetry (i.e., *r*_*b*_ = 1.0), promoting a well-defined modular arrangement. When asymmetry was introduced, by pruning reciprocal links and adding unidirectional ones, the cohesiveness of the network’s community structure diminished. In contrast, random networks with low initial reciprocity (i.e., *r*_*b*_ = 0.07) exhibited a notable enhancement in the modularity index as reciprocity increased.

The effect of varying reciprocity on network community structure shows class-dependent trends across weighted synthetic benchmarks and empirical connectomes (see Fig. B2). Networks initiated with high link or strength symmetry (e.g., modular, small-world, and human connectomes) exhibit comparatively modest changes in modularity with reciprocity and typically peak at intermediate values, sometimes plateauing or showing only a mild rollover at high reciprocity. By contrast, networks with initially asymmetric architectures (e.g., random graphs and the C. elegans connectome) display a monotonic increase in modularity across the examined reciprocity range. Taken together, these results indicate that reciprocity can progressively reinforce community organization in asymmetric networks, whereas in already symmetric networks the effect largely saturates beyond intermediate reciprocity

Community structures are crucial for the specificity and stability of real-world network topologies (Maslov and Sneppen, 2002). The NRC algorithms developed here systematically manipulate reciprocity, offering a valuable tool for examining how these structural features contribute to the overall dynamics and functionality of complex networks.

### 5.5 Clustering coefficient variations

The average clustering coefficients (Saramäki et al., 2007; Kaiser, 2008), detailed in Appendix Clustering coefficient, were examined across various networks by systematically varying reciprocity. The observed trends were strongly influenced by the networks’ initial reciprocity levels, indicating that the algorithm affects the relationship between reciprocity and clustering differently depending on the network configuration.

Changes in the average clustering coefficient with reciprocity closely track each network’s initial reciprocity (Fig. 4A). Small-world and modular benchmarks start at full link symmetry (*r*_*b*_ ≈ 1) and retain high clustering across the range, showing only modest decreases as symmetry is reduced by the algorithm. Human connectomes also begin near full reciprocity but exhibit a more noticeable decline in clustering as reciprocity decreases. In contrast, random graphs begin near (*r*_*b*_ ≈ 0) and display an approximately constant clustering coefficient across the reciprocity range, indicating that introducing reciprocity has little effect on local triadic density in the sparse random architectures. Other empirical connectomes (macaque, rat, mouse, Drosophila, C. elegans) with the initial binary reciprocity values ranging between 0.5 < *r*_*b*_ < 1.0 show mild, network-specific adjustments, typically small increases or near-stable behavior, consistent with their partial initial symmetry.

In weighted networks, both synthetic and empirical (Fig. B2), the average clustering coefficient is comparatively insensitive to reciprocity adjustments. Notable exceptions are the macaque, Drosophila, and C. elegans connectomes, which start at different initial reciprocity levels and span distinct weight scales; for these, both increases and decreases in reciprocity implemented by Algorithm 2 (weighted) are associated with reductions in the weighted clustering coefficient (Figs. 4 and B2). Importantly, the average clustering coefficient for weighted, directed networks is sensitive to the scale and distribution of edge weights; because we do not normalize weights across datasets, our plots are not intended for cross-network comparison of absolute magnitudes. Instead, we report within-network trajectories as reciprocity is varied.

## 6 Computational implications of graded reciprocity

The computational implications of graded reciprocity were explored through the memory capacity (Damicelli et al., 2022) of reservoir computing (RC) models. RC is a biologically inspired paradigm for processing temporal signals using recurrent neural networks (RNNs) (Jaeger and Haas, 2004; Maass et al., 2002). It consists of an input layer, a fixed recurrent reservoir, and a readout layer. The fixed-weight reservoir projects inputs into a high-dimensional dynamic state space, decoupling learning from structural adaptation. Only the readout layer is trained, which enables efficient exploration of how network topology, including reciprocity, affects computational capacity (Rodriguez et al., 2019; Damicelli et al., 2022; Suárez et al., 2020; Hadaeghi et al., 2025).

In our experiments, reservoir dynamics followed a leaky integrator (LI) model (Jaeger et al., 2007), where the leak rate *α* ∈ [0, 1] controlled temporal inertia: higher *α* produced faster responses, while lower values introduced slower dynamics. We set *α* = 1 and Δ*t* = 1, which improved memory capacity in our tests. The state update was:

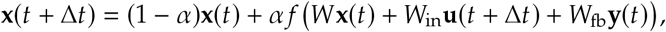

where *f* is tanh, and *W, W*_in_, *W*_fb_ denote the recurrent, input, and feedback weights.

The output was computed as:

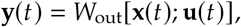

where *W*_out_ was trained via ordinary least squares regression with Tikhonov regularization:

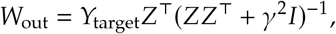

with γ = 10^−5^ and *Z* the concatenated reservoir and input states. The linear output activation ensured direct mapping from states to outputs.

The memory capacity of RC models was quantified using a delayed memory task. Each reservoir received a one-dimensional white noise input *u*(*t*) ~ 𝒩 (0, 1) and was trained to reconstruct delayed versions *d*_*k*_ (*t*) = *u*(*t* − *k*), with *k* = 1, 2, …, 1.4*N*_*x*_. MC was computed as

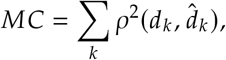

where *ρ*^2^ denotes the squared Pearson correlation coefficient on held-out data.

To investigate the effects of graded reciprocity, a series of experiments was conducted on binary and weighted random networks with adjusted reciprocity values. The performance of these networks was then assessed in a memory task, focusing on their ability to store and recall information. As shown in Figure 5, the influence of graded reciprocity on memory capacity (MC) was found to be noticeable in both binary and weighted networks, though the patterns observed differ. In binary networks, increasing reciprocity leads to a reduction in memory capacity, whereas in weighted networks, the opposite trend is observed, with memory capacity improving as reciprocity increases.

**Figure 5.**
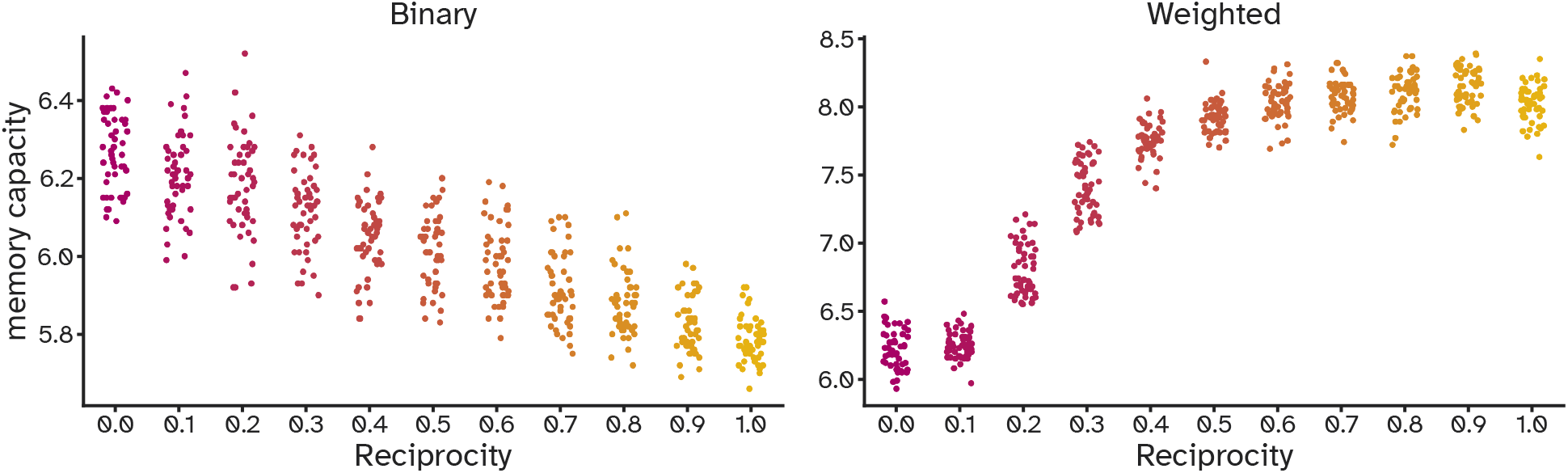
Trends in memory capacity of reservoir computing models with random network structures as their reciprocity is adjusted from *r* = 0 to *r* = 1.0 using Algorithm 1 (binary) and Algorithm 2 (weighted). To assess the memory of a given reservoir computing model, a sequence of white noise is generated by sampling from a normal distribution with mean 0 and variance 1.0 (i.e., *u* (*t*) ~ 𝒩 (0, 1)). Then *k* independent output units are trained to predict delayed versions of the input, i.e., *d*_*k*_ *t* = *u t k*. On the test dataset, all the predictions, *d*_*k*_ (*t*), are collected and the memory capacity (MC) is calculated using the squared Pearson correlation coefficient, *ρ*^2^ as 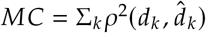. In the reservoir computing (RC) models used in this study, the hyperbolic tangent (tanh) function is employed as the transfer function, while a linear readout layer is used to map the reservoir states to the output. The input-to-reservoir weights are randomly sampled from a uniform distribution, ensuring a broad range of input connections. To satisfy the echo state property (ESP), the adjacency matrices are normalized such that their spectral radius is set to 1.0, thereby ensuring network stability and appropriate dynamics for computation over time (Jaeger, 2002).

In binary networks, increasing reciprocity and the resulting feedback loops between nodes reduce the overall diversity of the network’s states, limiting its ability to store and recall complex patterns. In fact, reciprocal connections tend to reinforce the same directions of information flow, potentially causing the recurrent network to become more deterministic and generating a poor repertoire of representations of the input stimulus. This limitation can undermine the ability of reservoir computing to create the diverse and rich representations necessary for effective computation.

On the other hand, increasing strength reciprocity leads to the modulation of a small subset of weights, causing them to have larger values while suppressing the smaller weights even further. As a result, increasing the reciprocity of a weighted network expands the range of weight values. While the total sum of the weights remains constant, their distribution shifts, resulting in a greater disparity among individual weights. This redistribution enhances the network’s memory capacity by making the system more selective in its interactions. The left panel of Figure 6 illustrates the average weight values at different levels of strength reciprocity, computed over 50 random matrix trials.

**Figure 6.**
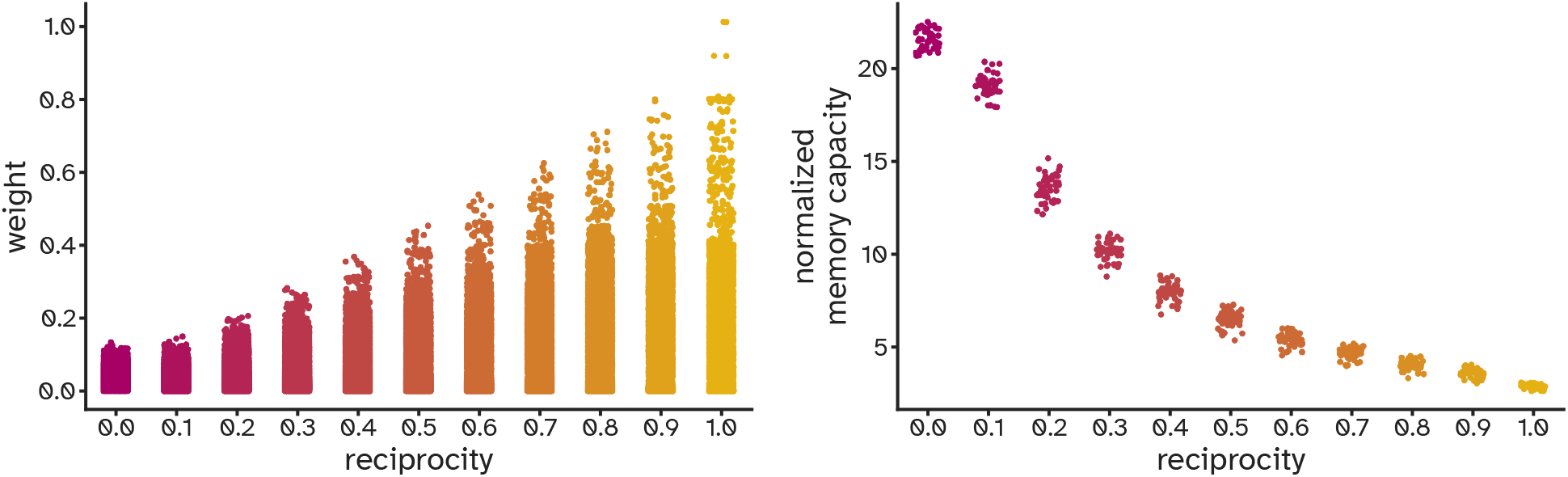
Weight modulation and its effect on memory capacity. Left: Average weight values at different levels of strength reciprocity, calculated over 50 random matrix trials. Right: Memory capacity of reservoir computing models with random network structures, as reciprocity is varied from *r* = 0 to *r* = 1.0 using Algorithm 2 (weighted), and subsequently normalized by dividing by the standard deviation of non-zero weights.

Since weight modulation introduces a confounding effect when analyzing memory capacity, MC was normalized through division by the standard deviation of weights. This normalization, as shown in the right panel of Figure 6, allows the impact of reciprocity on memory performance to be isolated. After normalization, the trend observed in the memory capacity closely resembles the one seen in the binary networks.

The experiments presented here form the basis for a more extensive analysis in our recent work, where we systematically applied these reciprocity control algorithms to explore memory capacity and the quality of internal representations in connectome-constrained reservoir computing models, with connection weights sampled from a log-normal distribution (Hadaeghi et al., 2025).

## 7 Performance and scalability

To evaluate the scalability of the reciprocity adjustment algorithm, tests were conducted on random networks with up to 2, 048 nodes and densities of 0.1, 0.2, 0.3, 0.4, and 0.5. Both binary and weighted networks were initialized with reciprocity values matching respective densities.

The binary algorithm consistently completed execution in under 1 second, with computation time primarily influenced by network size and the magnitude of reciprocity adjustment, as illustrated in Figure 7. The estimated time complexity of Algorithm 1 (binary) is 𝒪(*n*^2.28^), based on a curve fit of empirically measured runtimes as a function of network size over the tested range. This value reflects observed scaling behavior in our experiments rather than a formal analytic complexity bound. In contrast, the weighted network algorithm, described in Algorithm 2 (weighted), incorporates an iterative procedure that significantly affects performance. For 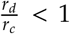, where *r*_*d*_ and *r*_*c*_ represent the desired and current reciprocity, the algorithm maintained high efficiency. However, when 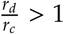, a time complexity of 𝒪 (*e*) was observed, reflecting the iterative nature of Algorithm 2 (weighted). This disparity in performance between binary and weighted networks highlights the added complexity introduced by weighted networks.

**Figure 7.**
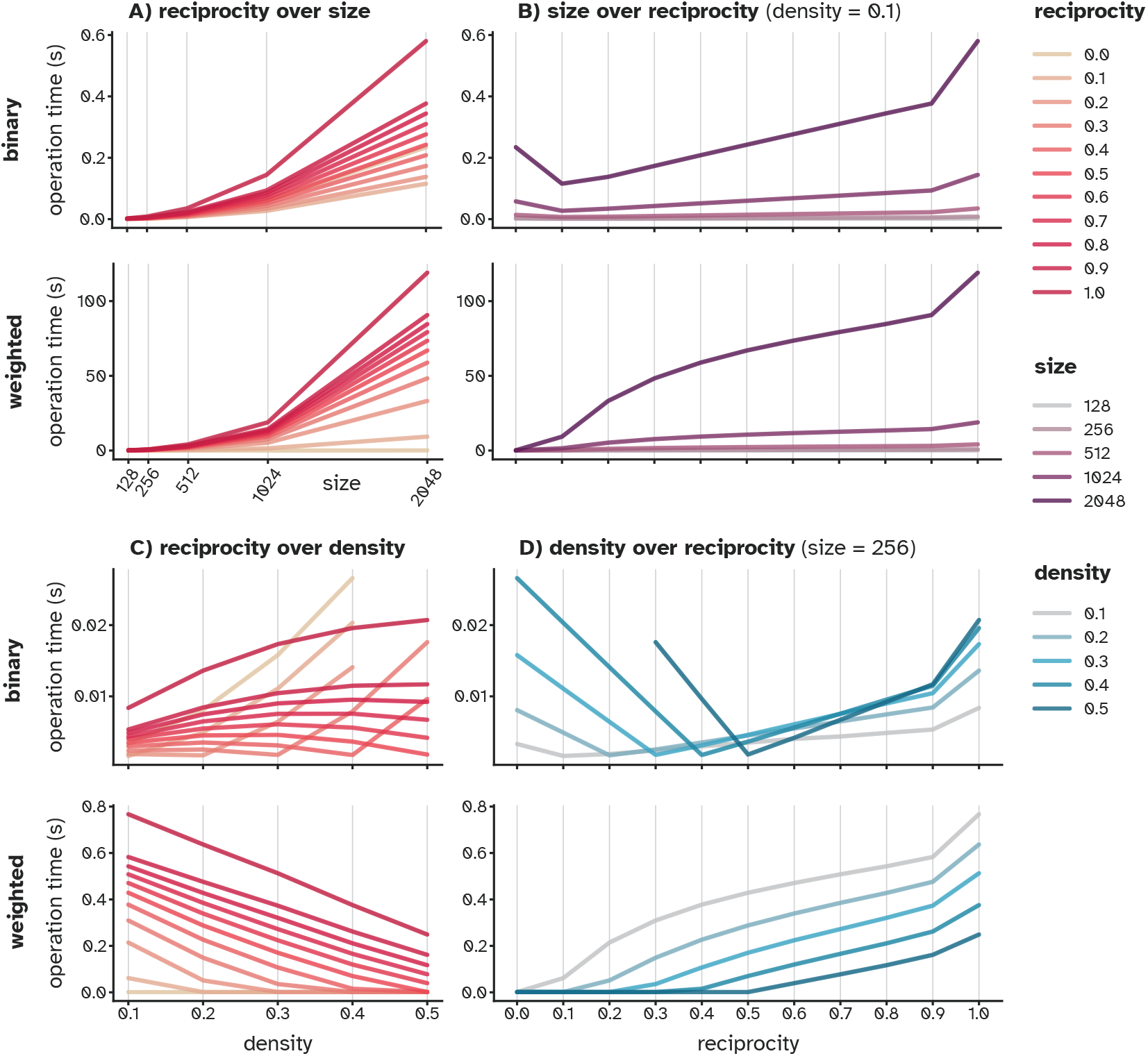
Scalability of Algorithm 1 (binary) and Algorithm 2 (weighted) as a function of network size, density, and initial link/strength reciprocity. A, B) Computation time for both algorithms primarily depends on the network size and the extent of reciprocity adjustment. The initial random network has low link/strength reciprocity (< 0.1), requiring the redistribution of unidirectional links in Algorithm 1 (binary) and an iterative procedure in Algorithm 2 (weighted). As network size increases, the number of links and elements requiring redistribution (Algorithm 1) or weight manipulation (Algorithm 2) grows, leading to longer computation times. C) In both binary and weighted networks initialized with reciprocity matching their density, this reciprocity(density) value represents a breakpoint in the piecewise linear trends of operation time. For a fixed number of nodes (*N* = 256), increasing density raises the number of links to be redistributed for higher reciprocity, thus increasing operation time in Algorithm 1 (binary). In weighted networks, higher density results in a greater number of non-zero elements in the adjacency matrix, allowing Algorithm 2 to operate on more values and converge faster. D) Computation time follows a piecewise relationship with the magnitude of reciprocity change. Breakpoints correspond to the original reciprocity, with segments on the left representing the time required to reduce reciprocity and those on the right showing the time needed to increase it in both algorithms.

## 8 Outlook

In complex systems, directed networks are prevalent, as they effectively capture the inherent asymmetry in both natural and artificial interactions. This prevalence is evident in representative datasets from the Netzschleuder network catalogue and repository (https://networks.skewed.de), where 92, 309 of 163, 735 networks are directed. From biological and neural networks to transportation and trade networks to social media, these networks rely on a delicate balance between reciprocity and asymmetry, which influences performance, adaptability, and resource utilization across various domains. Despite its significance, the role of graded reciprocity is often overlooked due to the challenge of modifying it without affecting other network properties.

We have developed novel network reciprocity control algorithms to adjust reciprocity in both binary and weighted directed networks while maintaining key characteristics such as density and total strength. This innovation provides a practical tool for exploring how reciprocity impacts network dynamics in fields such as complex systems physics, neuroscience, and bio-inspired computing.

While we focused here on topological and computational consequences, the NRC algorithms are readily applicable to future studies involving nonlinear dynamics, such as synchronization phenomena, phase transitions, critical cascades, and self-organized behavior in complex systems, where reciprocity plays a key regulatory role.

The approach also generates null models for more robust statistical analyses, particularly in computational neuroscience. In neuroscience, where symmetric connectome data may obscure underlying asymmetries (Suarez et al., 2022), our approach facilitates the exploration of synthetic asymmetric reciprocity, offering insights into the optimal balance for neural processing.

In artificial intelligence, our algorithm could enhance recurrent neural networks (RNNs), potentially improving training efficiency by fine-tuning the reciprocity-asymmetry balance. We demonstrated its broad applicability by examining how reciprocity influences network features, such as degree distributions, spectral properties, modularity, and clustering. The scalability of the NRC approach ensures its relevance to large-scale networks, providing deeper insights into the organization and dynamics of complex systems.

A key feature of the proposed NRC framework is its structural minimalism. By focusing exclusively on reciprocity adjustments while preserving connection density or strength, the method avoids complex rewiring schemes that may introduce unintended structural correlations. This simplicity makes the approach broadly applicable as a null model generator or control mechanism in network studies, and particularly useful for probing the specific role of directionality in nonlinear dynamics and information processing.

In sum, this work provides a tool for network hypothesis testing and exploration, offering new insights into understanding and optimizing binary and weighted directed network structures across various scientific fields.

## 9 Data and code availability

Data supporting the findings of this study are available publicly. The connectome dataset is available at netneurotools https://netneurotools.readthedocs.io. Synthetic data and codes that were used to implement the algorithm and perform network analysis in this manuscript are available via GitHub: https://github.com/m00rcheh/NRC_binary_and_weighted_Network_Reciprocity_Control.

## 10 Acknowledgments

The funding is gratefully acknowledged: F.H: DFG TRR169-A2, K.F: German Research Foundation (DFG)-SFB 936-178316478-A1; TRR169-A2; SPP 2041/GO 2888/2-2 and the Templeton World Charity Foundation, Inc. (funder DOI 501100011730) under grant TWCF-2022-30510. C.H: SFB 936-178316478-A1; TRR169-A2; SFB 1461/A4; SPP 1212 2041/HI 1286/7-1, the Human Brain Project, EU (SGA2, SGA3). We thank Arnaud Messé for valuable feedback and for reviewing the manuscript.

## Appendices

### Network properties

#### Non-normality

To quantify the departure from normality of the adjacency matrix, *W*, as its reciprocity is progressively reduced, we compute Henrici’s index. This measure evaluates the non-normality of a directed network by comparing its Frobenius norm, 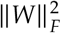, to its spectral norm:

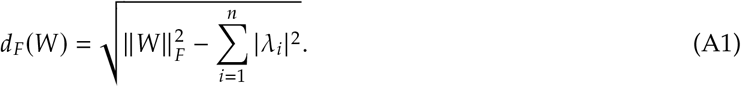

Here, 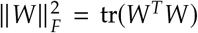 defines the Frobenius norm, while 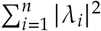 is the sum of squared eigenvalue moduli. For normal matrices, these are equal, but deviations quantify non-normality, as captured by Henrici’s index.

To enable comparison across systems of different sizes, the normalized index was suggested in (Asllani et al., 2018) as:

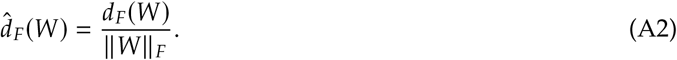

Figure A1 illustrates the normalized Henrici index as the reciprocity of synthetic benchmark networks is adjusted from *r* = 0 to *r* = 1 using Algorithm 1 (left panel) and Algorithm 2 (right panel).

#### Global efficiency

Global efficiency (Seguin et al., 2018; Latora and Marchiori, 2001) is a measure of how efficiently information is exchanged across a network. It quantifies the average inverse shortest path length between all pairs of nodes, reflecting the network’s overall connectivity and integration.

The formula for global efficiency *E*^∗^ is given by:

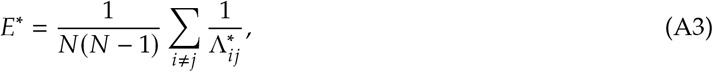

where *N* is the total number of nodes in the network, 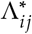. is the shortest path length between node *i* and node *j*, and the sum 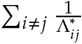 runs over all pairs of distinct nodes *i* and *j* in the network.

#### Modularity index

To quantify the strength of community structure in directed networks, we computed the modularity index *Q* using a formulation adapted from Leicht and Newman (Leicht and Newman, 2008). This version generalizes the well-known modularity measure for undirected networks to account for the directionality of edges. Specifically, the modularity is defined as:

**Figure A1.**
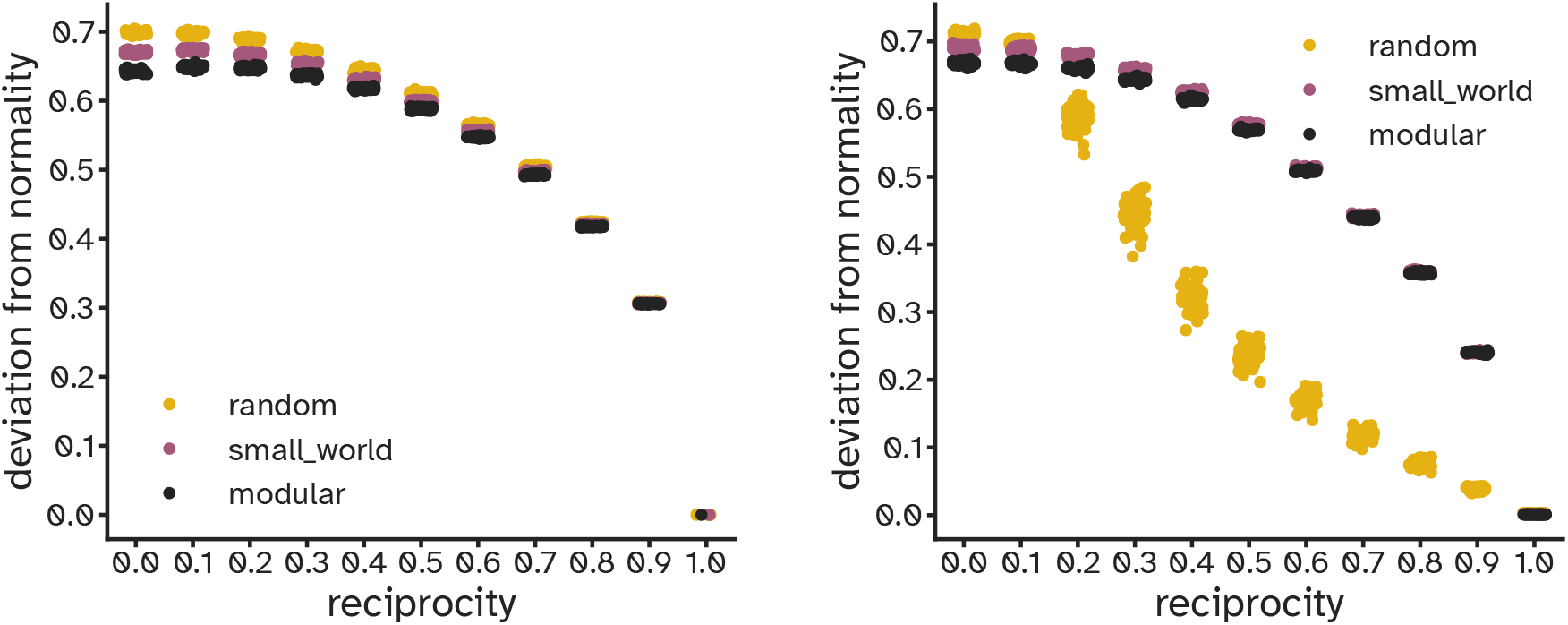
Functional implications of controlling reciprocity. Normalized index of departure from normality (Asllani et al., 2018) computed for 150 adjacency matrices (50 each for random, small-world, and modular structures) as their reciprocity is adjusted from *r* = 0 to *r* = 1.0 using Algorithm 1 (left panel) and Algorithm 2 (right panel). Across all cases, higher reciprocity consistently enhances normality, though different patterns emerge between random binary and weighted networks. Random networks, with initial reciprocity below 0.1, require iterative adjustments by Algorithm 2 (weighted) to reach higher target reciprocity values. In contrast, structured networks starting with *r* = 1.0 do not require iterative procedures, leading to distinct trends as reciprocity changes in Algorithm 2 (weighted).

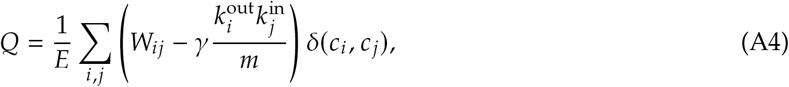

where *W*_*ij*_ is the weight of the directed edge from node *i* to node 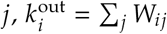 and 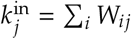 are the out- and in-strength of nodes respectively, 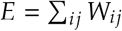 is the total edge weight, γ is a resolution parameter, and *δ* (*c*_*i*_, *c*_*j*_)is 1 if nodes *i* and *j* belong to the same community and 0 otherwise. We set γ = 1 in all our experiments.

To optimize *Q*, we employed a spectral bisection approach that recursively partitions the network using the leading eigenvector of the modularity matrix 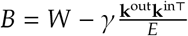. At each step, if the modularity gain is positive, the current module is split; otherwise, it is retained a^*E*^s a community. This procedure continues until no further partitioning improves the modularity. Community assignments were then used to compute the final modularity index. This method is deterministic and does not rely on stochastic optimization. We implemented this approach by slightly modifying the `modularity_dir` function provided by the bctpy library (Rubinov and Sporns, 2010).

#### Clustering coefficient

We computed the average clustering coefficient for both binary and weighted directed networks using methods provided by the bctpy library, which implements the approach introduced by Fagiolo (Fagiolo, 2007). For binary directed networks, we used the `clustering_coef_bd` function, which calculates the fraction of closed triads (3-node cycles) around each node. In directed graphs, each triad can be completed in up to eight possible ways due to edge directionality. The total number of such 3-cycles for a node is computed from the main diagonal of *S*^3^ / 2, where *S* = *A* + *A*^*T*^ is the symmetrized adjacency matrix. The maximum number of potential triangles accounts for both the node’s in-degree and out-degree, excluding mutual edge pairs that do not contribute to distinct triangle configurations. The final clustering coefficient is defined as the ratio of the number of observed triangles to the maximum number of possible triangles that can exist per node.

For weighted directed networks, we used the `clustering_coef_wd` function, which extends the binary formulation by incorporating edge weights. In this version, the weights matrix is transformed by taking the cube root of each edge weight to balance the influence of strong versus weak connections. The resulting symmetrized weighted matrix is then used to compute the weighted number of 3-cycles, again normalized by the same denominator used in the binary case. The mean clustering coefficient across nodes was used as a summary measure of local network organization.

### B Supplementary figures for weighted networks

**Figure B1.**
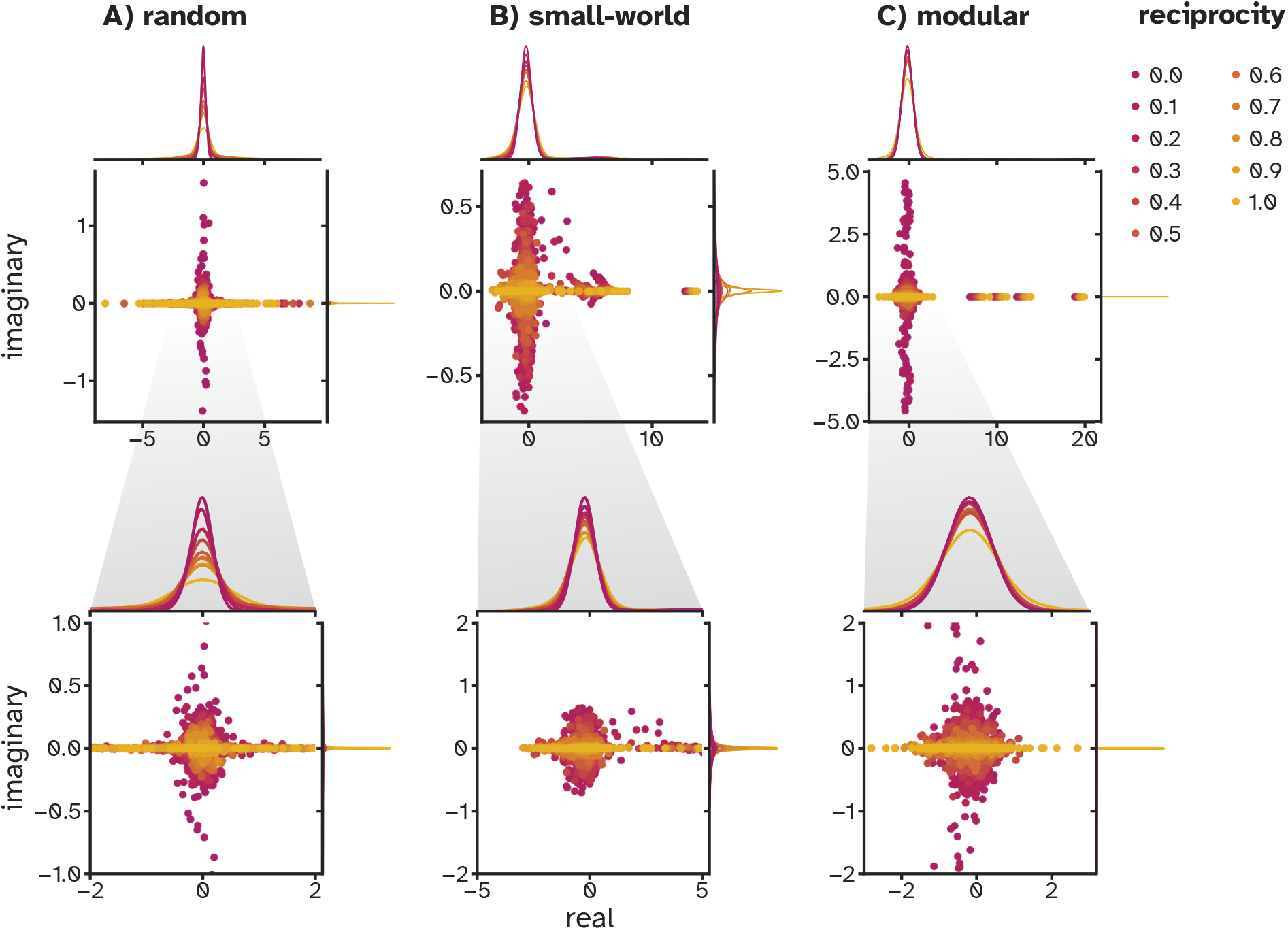
Evolution of eigenvalues of adjacency matrices as their reciprocity is adjusted from *r*_*b*_ = 0 to *r*_*b*_ = 1.0 using Algorithm 1 (binary). The top panel shows the original eigenspectrum, while the bottom panel provides a zoomed-in view. Distributions are calculated from 50 trials for each setting.

**Figure B2.**
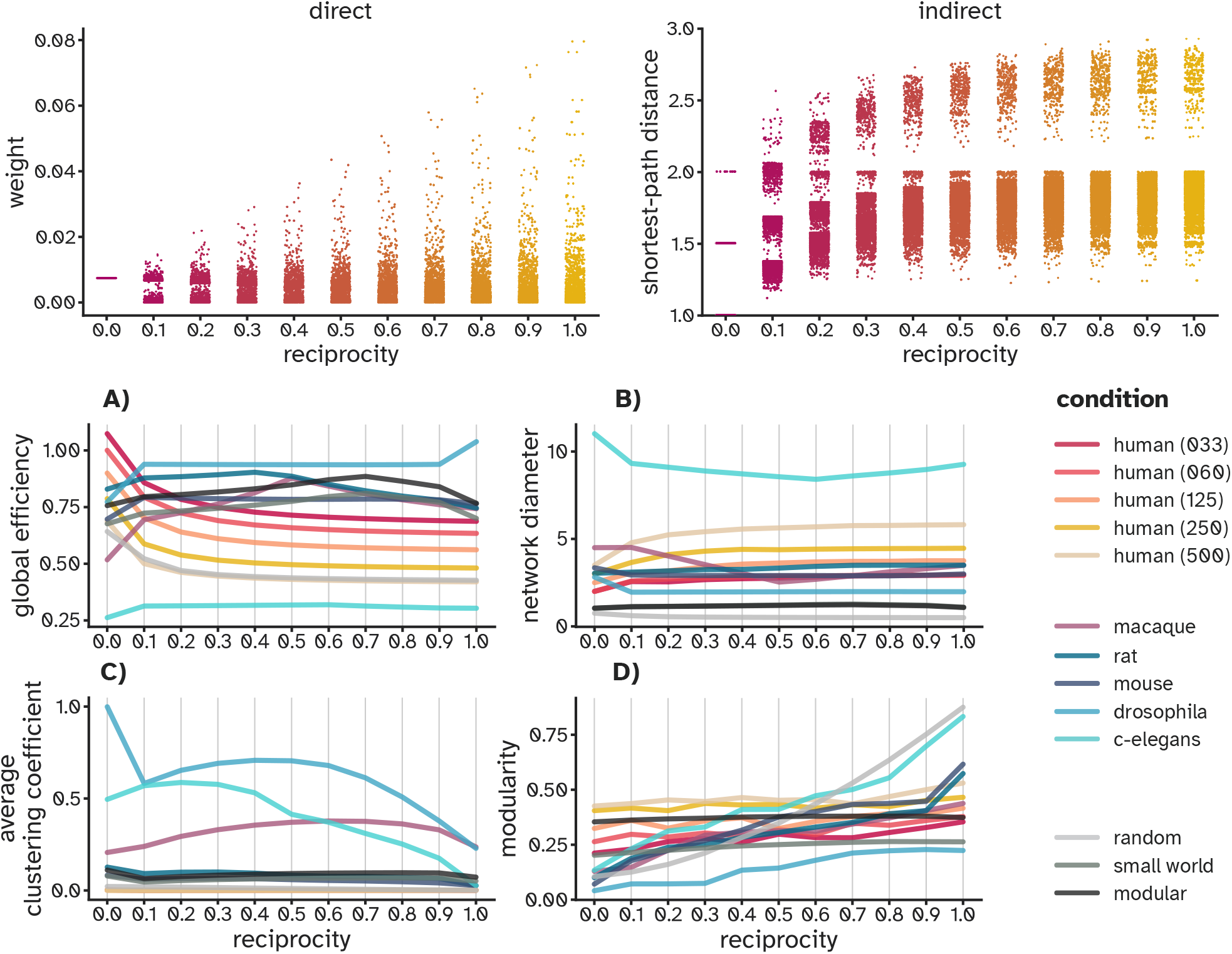
Structural implications of controlling strength reciprocity in weighted networks. Top panel: changes in weights and shortest paths in human connectome, denoted as ‘Human-struct-scale033’ in Table 1, as its reciprocity is adjusted from *r*_*w*_ = 1 to *r*_*w*_ = 0 using Algorithm 2 (weighted). Bottom panel: trends in key network metrics—global efficiency, network diameter, modularity index, and average clustering coefficient—in weighted adjacency matrices as their reciprocity is adjusted from *r* = 0 to *r* = 1.0 using Algorithm 2.

## Footnotes

1 Macaque connectome is only available as a binary network.

## References

Antoine Allard, M Ángeles Serrano, and Marián Boguñá. Geometric description of clustering in directed networks. Nature Physics, 20(1):150–156, 2024.

Malbor Asllani, Renaud Lambiotte, and Timoteo Carletti. Structure and dynamical behavior of non-normal networks. Science Advances, 4(12):eaau9403, 2018.

Vladimir Batagelj and Ulrik Brandes. Efficient generation of large random networks. Physical Review E—Statistical, Nonlinear, and Soft Matter Physics, 71(3):036113, 2005.

Marián Boguñá and M Ángeles Serrano. Generalized percolation in random directed networks. Physical Review E—Statistical, Nonlinear, and Soft Matter Physics, 72(1):016106, 2005.

Mihail Bota, Olaf Sporns, and Larry W Swanson. Architecture of the cerebral cortical association connectome underlying cognition. Proceedings of the National Academy of Sciences, 112(16):E2093–E2101, 2015.

Ann-Shyn Chiang, Chih-Yung Lin, Chao-Chun Chuang, Hsiu-Ming Chang, Chang-Huain Hsieh, Chang-Wei Yeh, Chi-Tin Shih, Jian-Jheng Wu, Guo-Tzau Wang, Yung-Chang Chen, et al. Three-dimensional reconstruction of brain-wide wiring networks in drosophila at single-cell resolution. Current Biology, 21(1): 1–11, 2011.

Aaron Clauset, Mark EJ Newman, and Cristopher Moore. Finding community structure in very large networks. Physical Review E—Statistical, Nonlinear, and Soft Matter Physics, 70(6):066111, 2004.

Fabrizio Damicelli, Claus C Hilgetag, and Alexandros Goulas. Brain connectivity meets reservoir computing. PLoS Computational Biology, 18(11):e1010639, 2022.

Giorgio Fagiolo. Clustering in complex directed networks. Physical Review E—Statistical, Nonlinear, and Soft Matter Physics, 76(2):026107, 2007.

Diego Garlaschelli and Maria I Loffredo. Patterns of link reciprocity in directed networks. Physical Review Letters, 93(26):268701, 2004.

Diego Garlaschelli and Maria I Loffredo. Multispecies grand-canonical models for networks with reciprocity. Physical Review E—Statistical, Nonlinear, and Soft Matter Physics, 73(1):015101, 2006.

Alessandra Griffa, Yasser Alemán-Gómez, and Patric Hagmann. Structural and functional connectome from 70 young healthy adults [data set]. Zenodo, 2019.

Fatemeh Hadaeghi, Kayson Fakhar, Moein Khajehnejad, and Claus Hilgetag. A computational perspective on the no-strong-loops principle in brain networks. bioRxiv, 2025. doi: 10.1101/2025.09.24.678310.

Joseph D Hart, Yuanzhao Zhang, Rajarshi Roy, and Adilson E Motter. Topological control of synchronization patterns: Trading symmetry for stability. Physical Review Letters, 122(5):058301, 2019.

Paul W Holland, Kathryn Blackmond Laskey, and Samuel Leinhardt. Stochastic blockmodels: First steps. Social Networks, 5(2):109–137, 1983.

Herbert Jaeger. Adaptive nonlinear system identification with echo state networks. Advances in Neural Information Processing Systems, 15, 2002.

Herbert Jaeger and Harald Haas. Harnessing nonlinearity: Predicting chaotic systems and saving energy in wireless communication. Science, 304(5667):78–80, 2004.

Herbert Jaeger, Mantas Lukoševičius, Dan Popovici, and Udo Siewert. Optimization and applications of echo state networks with leaky-integrator neurons. Neural Networks, 20(3):335–352, 2007.

Samuel Johnson. Digraphs are different: Why directionality matters in complex systems. Journal of Physics: Complexity, 1(1):015003, 2020.

Samuel Johnson, Virginia Domínguez-García, Luca Donetti, and Miguel A Munoz. Trophic coherence determines food-web stability. Proceedings of the National Academy of Sciences, 111(50):17923–17928, 2014.

Marcus Kaiser. Mean clustering coefficients: the role of isolated nodes and leafs on clustering measures for small-world networks. New Journal of Physics, 10(8):083042, 2008.

IA Kasyanov, Pim van der Hoorn, D Krioukov, and MV Tamm. Nearest-neighbor directed random hyperbolic graphs. Physical Review E, 108(5):054310, 2023.

Vito Latora and Massimo Marchiori. Efficient behavior of small-world networks. Physical Review Letters, 87 (19):198701, 2001.

Elizabeth A Leicht and Mark EJ Newman. Community structure in directed networks. Physical Review Letters, 100(11):118703, 2008.

Wenzhe Ma, Ala Trusina, Hana El-Samad, Wendell A Lim, and Chao Tang. Defining network topologies that can achieve biochemical adaptation. Cell, 138(4):760–773, 2009.

Wolfgang Maass, Thomas Natschläger, and Henry Markram. Real-time computing without stable states: A new framework for neural computation based on perturbations. Neural Computation, 14(11):2531–2560, 2002.

Natalia J Martinez, Maria C Ow, M Inmaculada Barrasa, Molly Hammell, Reynaldo Sequerra, Lynn Doucette-Stamm, Frederick P Roth, Victor R Ambros, and Albertha JM Walhout. A c. elegans genome-scale microrna network contains composite feedback motifs with high flux capacity. Genes & Development, 22(18):2535–2549, 2008.

Sergei Maslov and Kim Sneppen. Specificity and stability in topology of protein networks. Science, 296(5569): 910–913, 2002.

Everton S Medeiros, Ulrike Feudel, and Anna Zakharova. Asymmetry-induced order in multilayer networks. Physical Review E, 104(2):024302, 2021.

Dharmendra S Modha and Raghavendra Singh. Network architecture of the long-distance pathways in the macaque brain. Proceedings of the National Academy of Sciences, 107(30):13485–13490, 2010.

Takashi Nishikawa and Adilson E Motter. Symmetric states requiring system asymmetry. Physical Review Letters, 117(11):114101, 2016.

Sebastian Ocklenburg and Annakarina Mundorf. Symmetry and asymmetry in biological structures. Proceedings of the National Academy of Sciences, 119(28):e2204881119, 2022.

Louis M Pecora, Francesco Sorrentino, Aaron M Hagerstrom, Thomas E Murphy, and Rajarshi Roy. Cluster synchronization and isolated desynchronization in complex networks with symmetries. Nature Communications, 5(1):4079, 2014.

Nathaniel Rodriguez, Eduardo Izquierdo, and Yong-Yeol Ahn. Optimal modularity and memory capacity of neural reservoirs. Network Neuroscience, 3(2):551–566, 2019.

Mikail Rubinov and Olaf Sporns. Complex network measures of brain connectivity: uses and interpretations. Neuroimage, 52(3):1059–1069, 2010.

Mikail Rubinov, Rolf JF Ypma, Charles Watson, and Edward T Bullmore. Wiring cost and topological participation of the mouse brain connectome. Proceedings of the National Academy of Sciences, 112(32): 10032–10037, 2015.

Ueli Rutishauser, Jean-Jacques Slotine, and Rodney Douglas. Computation in dynamically bounded asymmetric systems. PLoS Computational Biology, 11(1):e1004039, 2015.

Jari Saramäki, Mikko Kivelä, Jukka-Pekka Onnela, Kimmo Kaski, and Janos Kertesz. Generalizations of the clustering coefficient to weighted complex networks. Physical Review E—Statistical, Nonlinear, and Soft Matter Physics, 75(2):027105, 2007.

Caio Seguin, Martijn P Van Den Heuvel, and Andrew Zalesky. Navigation of brain networks. Proceedings of the National Academy of Sciences, 115(24):6297–6302, 2018.

Tiziano Squartini, Francesco Picciolo, Franco Ruzzenenti, and Diego Garlaschelli. Reciprocity of weighted networks. Scientific Reports, 3(1):2729, 2013.

Laura E Suárez, Ross D Markello, Richard F Betzel, and Bratislav Misic. Linking structure and function in macroscale brain networks. Trends in Cognitive Sciences, 24(4):302–315, 2020.

Laura E Suarez, Yossi Yovel, Martijn P van den Heuvel, Olaf Sporns, Yaniv Assaf, Guillaume Lajoie, and Bratislav Misic. A connectomics-based taxonomy of mammals. eLife, 11:e78635, 2022.

Lav R Varshney, Beth L Chen, Eric Paniagua, David H Hall, and Dmitri B Chklovskii. Structural properties of the caenorhabditis elegans neuronal network. PLoS Computational Biology, 7(2):e1001066, 2011.

Duncan J Watts and Steven H Strogatz. Collective dynamics of ‘small-world’networks. Nature, 393(6684): 440–442, 1998.

Ronghua Xu, Qingpeng Zhang, and Sidie Tan. The formation of reciprocal social support in online support groups: a network modeling approach. IEEE Transactions on Computational Social Systems, 10(6):3370–3384, 2022.

Xing Yao, Yongzhong Zhang, Rizwana Yasmeen, and Zhen Cai. The impact of preferential trade agreements on bilateral trade: A structural gravity model analysis. PloS One, 16(3):e0249118, 2021.

Gorka Zamora-López, Changsong Zhou, Vinko Zlatić, and Jürgen Kurths. The generation of random directed networks with prescribed 1-node and 2-node degree correlations. Journal of Physics A: Mathematical and Theoretical, 41(22):224006, 2008a.

Gorka Zamora-López, Vinko Zlatić, Changsong Zhou, Hrvoje Štefančić, and Jürgen Kurths. Reciprocity of networks with degree correlations and arbitrary degree sequences. Physical Review E—Statistical, Nonlinear, and Soft Matter Physics, 77(1):016106, 2008b.

Yu-Xiao Zhu, Xiao-Guang Zhang, Gui-Quan Sun, Ming Tang, Tao Zhou, and Zi-Ke Zhang. Influence of reciprocal links in social networks. PloS One, 9(7):e103007, 2014.

Vinko Zlatić and Hrvoje Štefančić. Influence of reciprocal edges on degree distribution and degree correlations. Physical Review E—Statistical, Nonlinear, and Soft Matter Physics, 80(1):016117, 2009.

Vinko Zlatic and Hrvoje Štefancic. Model of wikipedia growth based on information exchange via reciprocal arcs. Europhysics Letters, 93(5):58005, 2011.

